# Evolution of Selective RNA Processing and Stabilization operons in cellulosome-harboring *Clostridium* spp

**DOI:** 10.1101/2021.06.12.447814

**Authors:** Yogendra Bhaskar, Mohammadhadi Heidari B., Chenggang Xu, Jian Xu

## Abstract

In selective RNA processing and stabilization (SRPS) operons, the stoichiometry of encoded proteins is determined by their respective 3’-end stem-loops (SLs), yet the evolution of this mechanism remains elusive. In cellulosomal operons of *Clostridium* spp., we show that the SLs and their associated genes form a monogamy companionship during the operon evolution. Based on ΔG of such SLs, we propose CoSLOE (Composite SL-based Operon Evolution) model with evolutionary ratio (ER) >1 or <1 for positive or negative selection of SRPS operons. In the composite SL-ΔG-based tree (CoSL-tree) of cellulosomal operons, when traversing from leafs to the root nodes, ERs reveal diversifying/positive selection towards a less efficient cellulosomal system, consistent with glycoside-hydrolase gene variation both in-operon and genome-wide. A similar pattern is followed by the ATPase operon and the majority of orthologous SRPS operons genome-wide, suggesting conservation among operons in such selection. Thus SRPS operons via their transcript-stabilizing non-coding elements are highlighting a link between operon stoichiometry and operon evolution.

## 1 Introduction

In bacterial genomes, ~50% of the genes are organized and regulated in the form of operon (Osbourn & Field *et al.*, 2009). Within an operon, to ensure proper absolute and relative abundance of the component genes, one strategy adopted by certain bacteria is selective RNA processing and stabilization (SRPS), where the RNA molecule is cleaved by ribonuclease into fragments, and then with the involvement of the specific *cis*-elements (Stem-loops), mature mRNA transcripts stabilize to differential gene expression and eventually to the protein complex (Rochat *et al.*, 2013). The SRPS mechanism controls operons that encode a variety of key protein complexes and regulatory pathways such as the glycolysis pathway, maltose transport system, cellulosome complex and photosynthetic apparatus (Newbury *et al.*, 1987, Klug *et al.*, 1993, Ludwig *et al.*, 2001, Xu *et al.*, 2015).

Using the cellulosome-encoding *cip-cel* operon of *Clostridum cellulolyticum* (*Ccel*) as a model, we showed that the stem-loops generally located at the 3’-end of regulated genes precisely regulate structure and relative abundance of the subunit-encoding transcripts processed from a primary polycistronic RNA (Xu *et al.*, 2015). Importantly, the “ratio” of subunit-encoding transcripts for the *cip-cel* operon, which quantitatively specifies cellulosome stoichiometry, appears to be encoded by the genome (i.e., organism-specific) and insensitive to alteration of culture conditions, since change among glucose, cellobiose and cellulose did not result in ratio change (Xu *et al.*, 2015). These findings revealed a key role of such stem-loops (SLs; i.e., all such SLs present in a SRPS operon) in specifying proper function of SRPS-operon-encoded protein complexes (or metabolic pathways). Moreover, they strongly suggest potential links between the structure and function of these stem loops to organismal evolution.

However, key questions remain unanswered: (*i*) how do these SLs evolve? How conserved are these SLs among orthologous operons? What is the nature of such conservation? (*ii*) What is the link in evolution between these SLs and their companion genes in SRPS operons? (*iii*) What roles do these SLs play in the evolution of SRPS operons? Are these roles conserved for SPRS operons at a genome-wide scale? How similar or divergent are these roles across different genomes? Can evolution of SRPS operons be quantitatively modeled via these SLs? Here in cellulosomal operons of *Clostridium* spp., based on ΔG of such SLs, we propose CoSLOE (Composite SL-based Operon Evolution) with evolutionary ratio (ER) >1 or <1 for positive or negative selection of SRPS operons. In the CoSL-tree of cellulosomal operons, when traversing from leafs to the root nodes, ERs reveal diversifying/positive selection towards a less efficient cellulosomal system, consistent with glycoside-hydrolase gene variation both in-operon and genome-wide. A similar pattern is followed by the ATPase operon and the majority of orthologous SRPS operons genome-wide, suggesting conservation among operons in such selection. Thus SRPS operons via their transcript-stabilizing non-coding elements are highlighting a link between operon stoichiometry and operon evolution.

## 2 Materials and Methods

### 2.1 Prediction of the stable stem-loops

SLs in *C. cellulolyticum* were predicted via the following steps (Bhaskar *et al.*, 2021). (*i*) Prediction of motifs using RNAmotif (Macke *et al.*, 2001); (*ii*) Estimation of free-energy and RNA secondary structure using RNAfold (Hofacker, 2003); (*iii*) Genome mapping of the predicted SLs; (*iv*) Screening of the SLs based on their stability for highly stable SLs. These stable SLs were then mapped to the operon map of the respective species, followed by functional classification based on the derived classification rules, whereby the SRPS operons were identified. Genome-encoded ratios were predicted for these SRPS operons using the ΔG of the harbored SLs (Bhaskar *et al.*, 2021).

### 2.2 Calculation of the ΔG-based ratio for an SRPS operon

Ratios were calculated in the SRPS operon using the ΔG (free-energy) of the SLs present in and flanking the operon (**Fig. 1A**). For example, the ratio for a four-gene operon (with SLs found after first two genes and at the end of operon) “Gene-1 (ΔG1): Gene-2 (ΔG2): Gene-3: Gene-4 (ΔG4)” would be “ΔG1: ΔG2: ΔG4: ΔG4”. To normalize the ratio, ΔG of all SLs in an operon were divided by the sum of all ΔG (**Table S1**).

**Figure 1.**
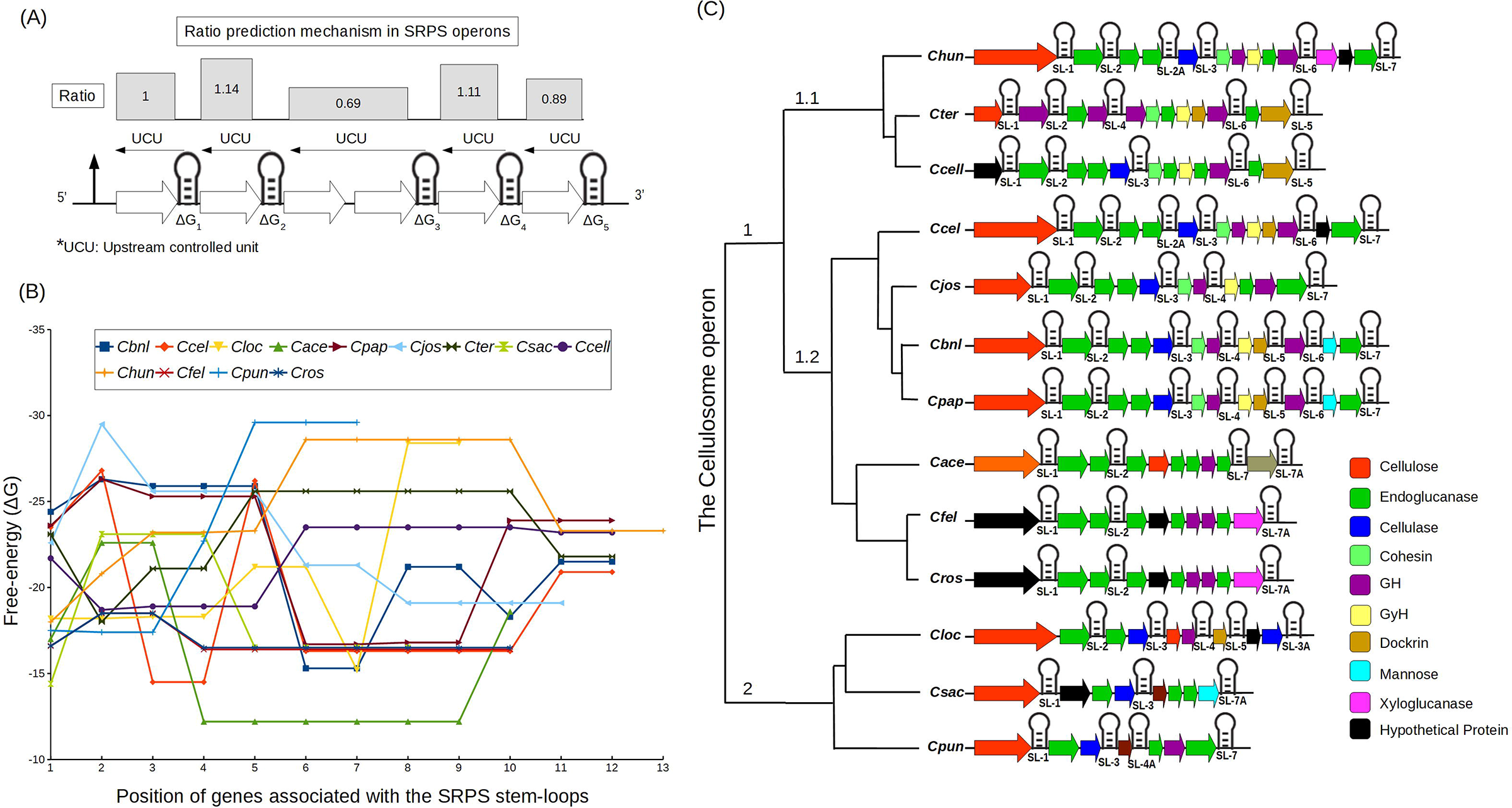
Composite SLs (CoSLs) in the cellulosomal operons from 13 Clostridial species. (**A**) Schematic representation of the SRPS operon via ΔG of the harbored composite SLs. Upstream Controlled Unit (UCU) represents the region (which is upstream to a SL and can include multiple genes) that is regulated by a SL via the SRPS mechanism. (**B**) ΔG of the harbored SLs in the cellulosomal operons from 13 *Clostridium* spp., showing skewness of the ΔG within an operon and divergence of pattern among orthologous operons. (**C**) Composite SLs (CoSLs)-based tree of the cellulosomal operons using the orthologous SLs. Genes are colored based on the encoded protein.

### 2.3 Phylogenetic analysis of the cellulosomal and the ATP synthase operons

The genomes and associated annotations of 13 cellulosome operon-harboring Clostridial species including *Ruminiclostridium cellulolyticum* H10 (*Ccel*; NC_011898.1), *Ruminiclostridium papyrosolvens* DSM 2782 (*Cpap*; GCF_000175795.2)*, Clostridium saccharoperbutylacetonicum* (*Csac*; NC_020291.1)*, Clostridium* sp. BNL1100 (*Cbnl*; GCF_000244875.1)*, Clostridium felsineum* DSM 794 (*Cfel*; GCF_002006355.1)*, Ruminiclostridium josui* JCM 17888 (*Cjos*; GCF_000526495.1)*, Ruminiclostridium cellobioparum* subsp. termitidis CT1112 (*Cter*; GCF_000350485.1)*, Clostridium acetobutylicum* ATCC 824 (*Cace*; NC_015687.1)*, Clostridium cellulovorans* 743B (*Cloc*; NC_014393.1)*, Ruminiclostridium hungatei* DSM 14427 (*Chun*; GCF_002051585.1)*, Clostridium puniceum* DSM 2619 (*Cpun*; GCF_002006345.1)*, Clostridium roseum* DSM 7320 (*Cros*; GCF_002006215.1) and *Ruminiclostridium cellobioparum* DSM 1351=ATCC 15832 (*Ccell*; GCF_000621505.1) (**Table S1**), were downloaded from NCBI. Cellulosome operons from *Cpap, Csac, Cbnl, Cfel, Cjos, Cter, Cace, Cloc, Chun, Cpun, Cros* and *Ccell* were identified by the available annotation and BLAST (Altschul *et al.*, 1990), where the *cip-cel* operon (encoding the cellulosome) from *Ccel* was used as a query with the e-value cutoff of 1e-5. Organismal phylogeny (16S-tree) of these species was derived using the 16S rRNA sequence, where all positions containing gaps and missing data were eliminated, which resulted in a total of 1,326 positions in the final multiple-sequence alignment. Phylogenetic analyses for the cellulosome and the ATPase operons were conducted in MEGA7 (Kumar *et al.*, 2016) via the Maximum Likelihood method.

The ΔG-based dendrogram of SLs was performed using the *pvclust* (Suzuki & Shimodaira, 2006) package in R (CRAN http://cran.r-project.org/) (**Fig. S1A**). To calculate the SLs’ ΔG-based dendrogram (CoSL-tree), ΔG of all SLs in an operon were divided by the sum of all ΔG, which generated a normalized proportion for an operon, and empty cells (i.e., values are “non-applicable”) were replaced by the average value of that proportion while clustering (**Table S1**). The normalized ΔG proportions of 13 clostridia species were supplied to *pvclust* with the Euclidean distance method and the *ward.D2* hierarchical clustering, with bootstrapping for 1000 times. The Ka/Ks values for genes were calculated using the Codeml tool of *PAML* package (Yang *et al.*, 2007).

### 2.4 Structural alignment analysis of orthologous SLs

The orthologous SLs from the cellulosomal operons and the ATP synthase operons were aligned structurally and sequence-wise using the *LocARNA* alignment and folding tool (Smith *et al.*, 2010, Will *et al.*, 2012). Evolution of the orthologous SLs was shown using the multiple-alignment of SL sequences and dot-bracket notations.

### 2.5 Derivation of Composite Stem-Loop-based Operon Evolution (CoSLOE) model

The CoSLOE model was described via an equation that calculates the evolutionary ratio (ER):

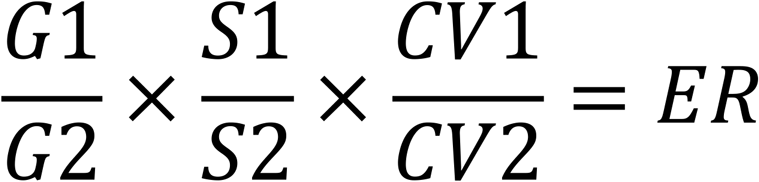

where G1 and G2, S1 and S2, CV1 and CV2 are the number of genes, number of SLs and coefficient of variations (CV) respectively, in the two operons. CV is the ratio of standard deviation (RatioSD) and mean 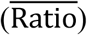 of the ΔG-based ratio of an operon.

## 3 Results

### 3.1 Phylogenetic analysis of the SLs in cellulosome-encoding SRPS operons from 13 Clostridial genomes

To probe the role of SLs in the function and evolution of SRPS operons at the whole-genome scale, we developed an approach to identify the SRPS operons based on the genome-wide predicted stable stem-loops (SLs) and then use the free-energy (ΔG) of these stable SLs to calculate ratios of SRPS operons (Bhaskar *et al.*, 2021) (**Fig. 1A**). The ΔG-based ratios were calculated for the cellulosome complex operon (cip-cel) in *Ccel*, which can model stoichiometry of the encoded complex. To probe how this mechanism has evolved, we extended the analysis to twelve additional mesophilic Clostridial spp.: *C. papyrosolvens* (*Cpap*)*, C. saccharoperbutylacetonicum* (*Csac*)*, C.* sp. BNL1100 (*Cbnl*)*, C. felsineum (Cfel), C. josui (Cjos), C. termitidis (Cter), C. acetobutylicum (Cace), C. cellulovorans (Cloc), C. hungatei (Chun), C. puniceum (Cpun), C. roseum (Cros),* and *C. cellobioparum* (*Ccell*; **Table S1**). These operons are orthologous, as indicated by orthology of genes, functional conservation of encoded proteins and the global similarity in operon structure. Our ΔG-based method predicted 7, 7, 5, 5, 5, 6, 5, 4, 3, 3, 3 and 3 SLs in *Cbnl, Cpap, Cjos, Cter, Ccell, Chun, Cloc, Cace, Cpun, Cros,* and *Cfel* respectively (**Table S1**). The ΔG-based ratio for these Clostridial species were also highly skewed, e.g., the ratios of *Cbnl* and *Cpap* are “-24.4:-26.3:-25.9:-25.9:-25.9:-15.3:-15.3:-21.2:-21.2:-18.3:-21.5:-21.5” and “-23.6:-26.3: - 25.3:-25.3:-25.3:-16.7:-16.7:-16.8:-16.8:-23.5:-23.9:-23.9” respectively. Similarly, *Ccell* and *Cter* exhibit identical ratios, so do *Cace*, *Cfel* and *Cros* (**Fig. 1B**; **Table S1**).

To probe how such operon properties have evolved, the ΔG-based proportions of all harbored SLs in an operon (which we termed “composite SLs” or CoSL) were used to generate a dendrogram (CoSL-tree; **Fig. S1A**). CoSL-tree was then compared to the 16S rRNA-based tree (16S-tree; i.e., the organismal phylogeny; **Fig. S1B**). Predicted ratios from the 13 cellulosomal operons were combined to form a data matrix, which was then used for the hierarchical clustering with 1000 iterations to generate the ratio-based tree (**Methods**). Intriguingly, the species were classified differently in the two clades derived from CoSL-tree (**Fig. S1A**) and 16S-tree (**Fig. S1B**). For example, (*i*) *Cace* and *Cros* are in Clade 1 of CoSL-tree, yet found in Clade 2 of 16S-tree; (*ii*) *Cpun* is an out-group in CoSL-tree, whilst *Cfel* is an out-group in 16S-tree; (*iii*) *Cros* and *Cfel* are clustered in CoSL-tree yet distantly apart in 16S-tree. Such difference between CoSL-tree and 16S-tree indicates the deviation of SRPS operon evolution from organismal taxonomy.

### 3.2 Gene-SL relationship during evolution of Clostridial cellulosomal operons

To probe the roles of SLs in cellulosomal operon evolution, seven orthologous SLs were first identified in the intergenic regions of the 13 orthologous cellulosomal operons, via comparison of their sequences, structures and organization in the operons (**Fig. 1C; Fig. 2**; **Fig. S2**). However, not all the Clostridial species harbor similar numbers of orthologous SLs and at identical positions (**Fig. 1C**; **Table S1**): 7 SLs in *Cbnl* and *Cpap*, 6 in *Ccel* and *Chun*, 5 in *Cjos, Cter, Ccell* and *Cloc*, 4 in *Cpun* and *Cace* and 3 in *Cfel*, *Cros* and *Csac*. The presence of these SLs suggests SPRS mechanisms in these 13 cellulosomal operons (for *Cloc*, the role of multiple promoters is also involved (Doi *et al.*, 1998)).

**Figure 2.**
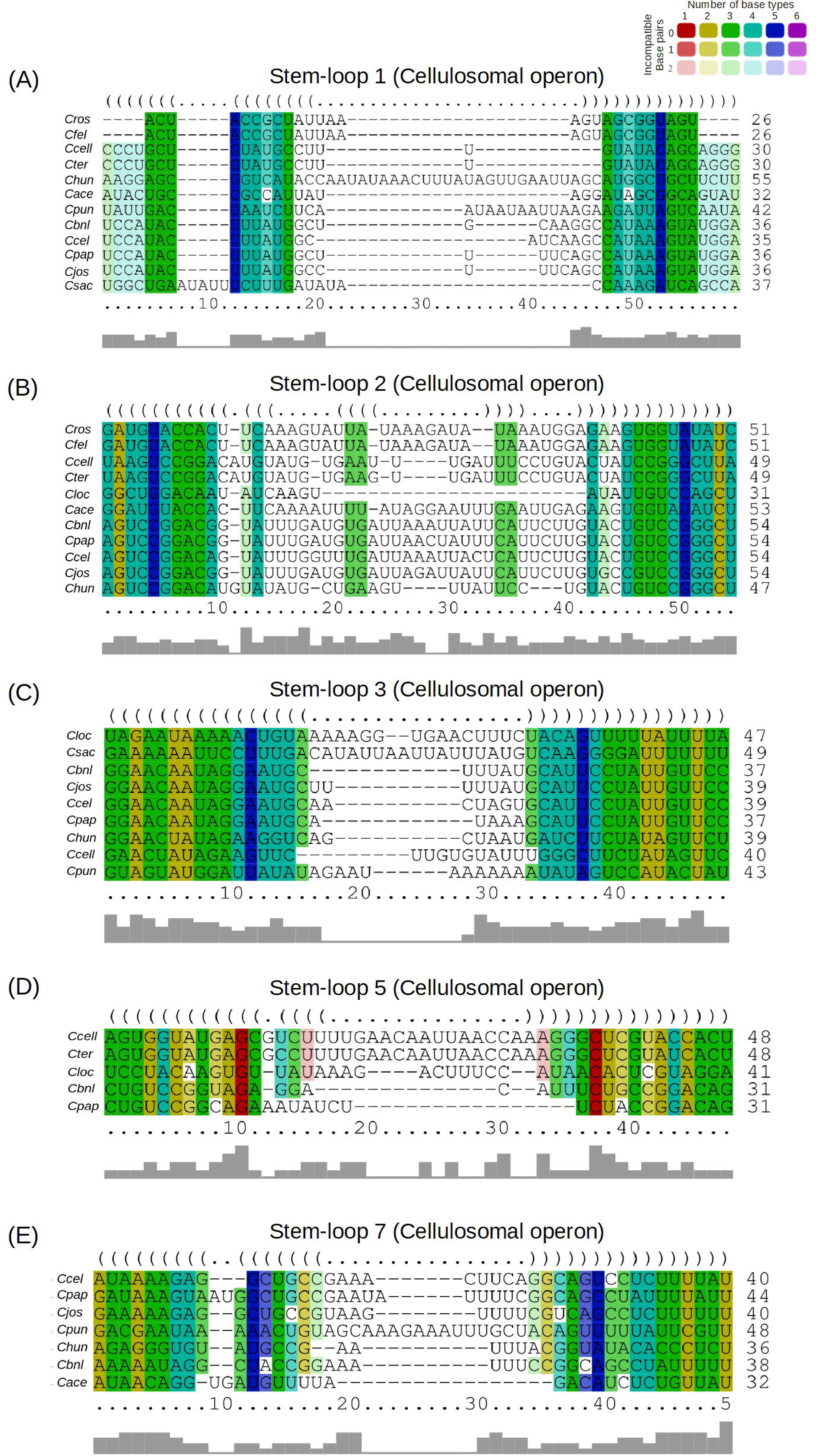
Structural alignment of SLs harbored in the cellulosome operons from 13 Clostridial species reveal the relationship between SLs and their associated genes. The multiple sequence alignment is shown for the five orthologous SLs: SL-1 (**A**), SL-2 (**B**), SL-3 (**C**), SL-5 (**D**) and SL-7 (**E**). SLs were aligned via sequence or structure using *LocARNA* and shown with the consensus structure in the Dot bracket form (middle). Compatible base pairs are colored based on the standard format, where the hue shows sequence conservation among the number of different types of compatible base pairs (C-G, G-C, A-U, U-A, G-U or U-G) in the corresponding columns. Color saturation decreases with the number of incompatible base pairs. The bar plot represents conservation of compatible base pairs (higher bar for higher conservation, and vice versa).

Interestingly, although the region between a SL and its associated genes can be inserted by another gene, the SLs are always positioned with their associated genes in a sequential fashion that is conserved among a set of orthologous operons. Thus, to probe the gene–SL relationship, orthologous SLs were aligned via sequence and structural similarity (**Fig. S2**). Compatible base pairs (in the stem sequences) were found in SL-1, 2, 3, 5 and 7, underscoring the structural similarity among the orthologous SLs (**Fig. 2**). Specifically, (*i*) SL-1 is present in all the Clostridial species (except *Cloc*), and SL-1 and 2, in their respective clades, are of similar length and identical ΔG to other orthologous SLs, yet show higher variation in their loop sequences (**Fig. 2A**, **B**; **Fig. S2A**, **B**); (*ii*) SL-3 shows less sequence variation in the two clades than SL-1 and 2, possibly due to its role as terminator SLs (**Fig. 2C**; **Fig. S2C**); (*iii*) SL-4, 5 and 6 are clade-specific, as they are absent in Clade 2 except the SL-4 in *Cloc* (**Fig. 2D**; **Fig. S2D**, **E**, **F**); (*iv*) similar to SL-3, SL-7 carries a low level of sequence variation (**Fig. 2E**; **Fig. S2G**). Such variation in SL sequence and structure depicts their evolutionary distance.

Intriguingly, a dockerin-encoding gene, located at the 7^th^ position of operon in *Cloc*, the 9^th^ in subclade of *Ccel-Cjos-Cbnl-Cpap* (except *Cjos*) and the 12^th^ in Subclade 1.1 species (except *Chun*) is always controlled by the orthologous SL-5 (**Fig. 1C, 2D**; **Fig. S2E**). Similarly, a cellulase-encoding gene, situated at distinct positions among cellulosomal operons, is controlled by an orthologous SL-3. In addition, clade-specific derivative homologous SLs in the cellulosomal operons also show such loyalty with their respective companion genes, e.g., (*i*) *Cloc* harbors an extra cellulase-encoding gene carrying SL-3A (homologous to SL-3; **Fig. S2H**); (*ii*) SL-2A is found in *Ccel* and *Chun*, similar to SL-2 (**Fig. S2I**); (*iii*) SL-7A is found in *Csac*, which is similar to SL-7 and works as terminator to the operon (**Fig. S2J**). These observations suggest monogamy as one feature of the gene-SL relationship during evolution of SRPS operons.

### 3.3 The Composite Stem-Loop based Operon Evolution (CoSLOE) model for SPRS operons

Taking advantage of the link between SLs and evolution of operon, we propose Composite Stem-Loop based Operon Evolution (CoSLOE) for the SRPS operons (**Fig. 3**). The model consists of (**Equation I**): (*i*) the number of genes in the operon (G), where the addition of one gene shows the positive selection, while an equal number of genes suggests neutral operons; (*ii*) the number of SLs (S), which plays crucial roles in regulation, stabilization and termination of genes; (*iii*) variance of ΔG of the SLs in operons (CV), where multiple SLs with distinct free-energy together specify and control the stoichiometry of gene expression. Therefore, the evolutionary ratio (ER) of an operon with respect to the other operons is,

**Figure 3.**
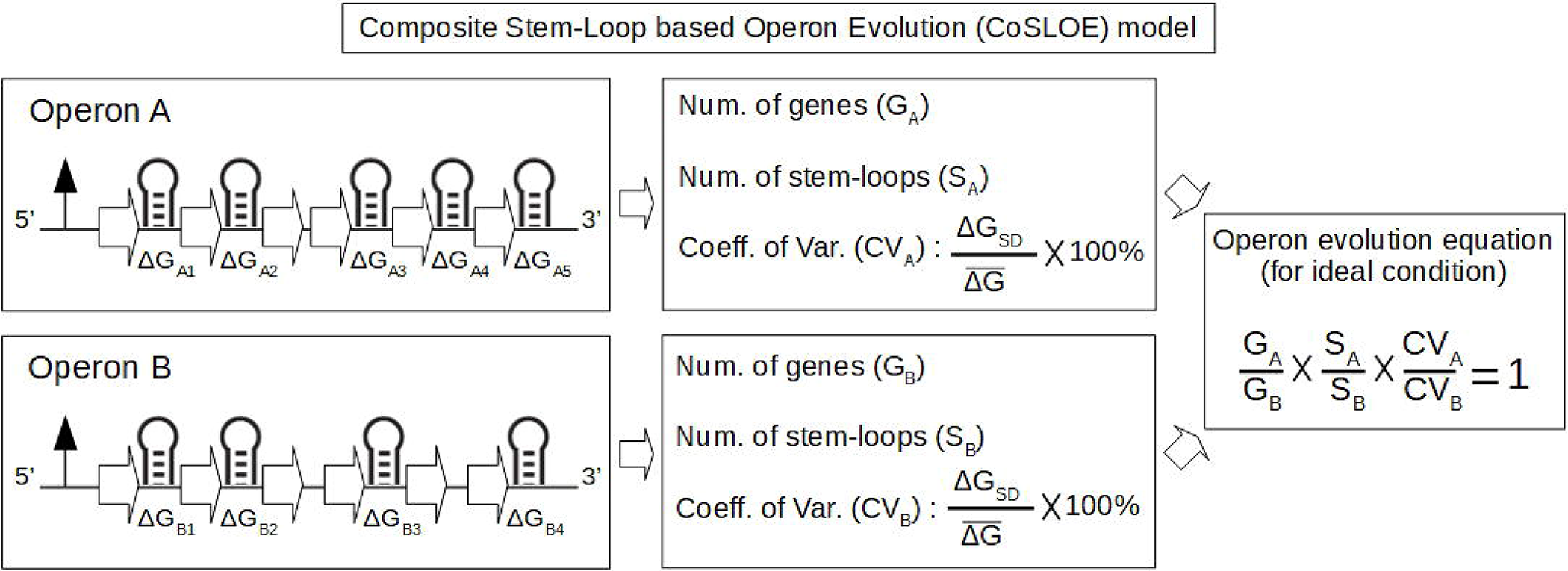
Proposing CoSLOE to quantitatively model the evolution of SRPS operons. In CoSLOE (Composite Stem-Loop based Operon Evolution), operon ER is calculated based on number of genes, number of SLs and coefficient of the variation (CV) of the ΔG-based (i.e., ΔG of the CoSLs) ratio. ΔG_SD_ and 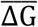 represent the standard deviation and the mean, respectively. The ratio determines the selection pressure between two operons, *i.e.,* ratio of 1 represents the neutral selection, while positive (or negative) selection occurs when ratio is >1 (or <1). These ratios in a clade of a tree determine the direction of the evolution, *i.e.* positive selection for species with ratio >1 and negative/purifying selection for species with ratio <1.

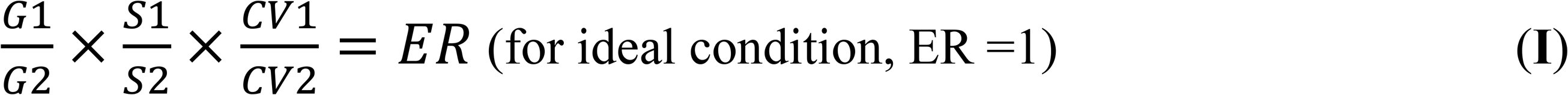

where G1 and G2, S1 and S2, CV1 and CV2 are the number of genes, number of SLs and coefficient of variations (CV) respectively, in the two operons. CV is ratio of standard deviation and mean of the ratio of ΔG of SLs for an operon. Positive or purifying selection of the operon is indicated by ER > 1 and ER < 1 respectively, while ER of 1 corresponds to neutral selection (i.e., ideal condition).

### 3.4 CoSLOE reveals selection pressure on the cellulosomal operons

To probe their evolution, pairwise ERs for the 13 cellulosomal operons in CoSL-tree were derived via CoSLOE (**Equation I**; **Fig. 3**). In Subclade 1.2 (**Fig. 1C**), (*i*) *Cbnl* and *Cpap* show ER of 1.01 and 0.99 with each other respectively, suggesting that the selection pressure is almost neutral and *Cbnl* is positively selected towards the root; (*ii*) the next nearest species is *Cjos*, which lacks one gene and two SLs possibly due to the deletion or horizontal transfer of genes, shows the ER of 0.57, 0.57 and 0.52 with *Cpap, Cbnl* and *Ccel* respectively (**Table 1A**), *i.e.* equally separated from all the three clostridia; (*iii*) however, the Ka/Ks values, at the gene level selection, for the first gene of *Cjos* are 1.42, 1.45, and 1.5 with *Ccel*, *Cbnl*, *Cpap* respectively (**Table S2**), suggesting the first gene of these operons is under positive selection towards *Cjos*; (*iv*) the operon ER for *Ccel* is 1.93 1.10 and 1.10 with *Cjos, Cbnl* and *Cpap* respectively (**Table 1A**), which depict the positive selection with the addition of a new gene at 11^th^ place in operon (**Fig. S5**). Taken together, in Subclade 1.2 of CoSL-tree, species are under positive selection while going from *Cpap* to *Ccel* (**Fig. 1C**), and also while going from *Cros* to *Cace* (due to the much higher *Cace-Cros* ER of 7.76 than *Cfel*-*Cros* ER of 1.05; **Table 1B**).

**Table 1.**
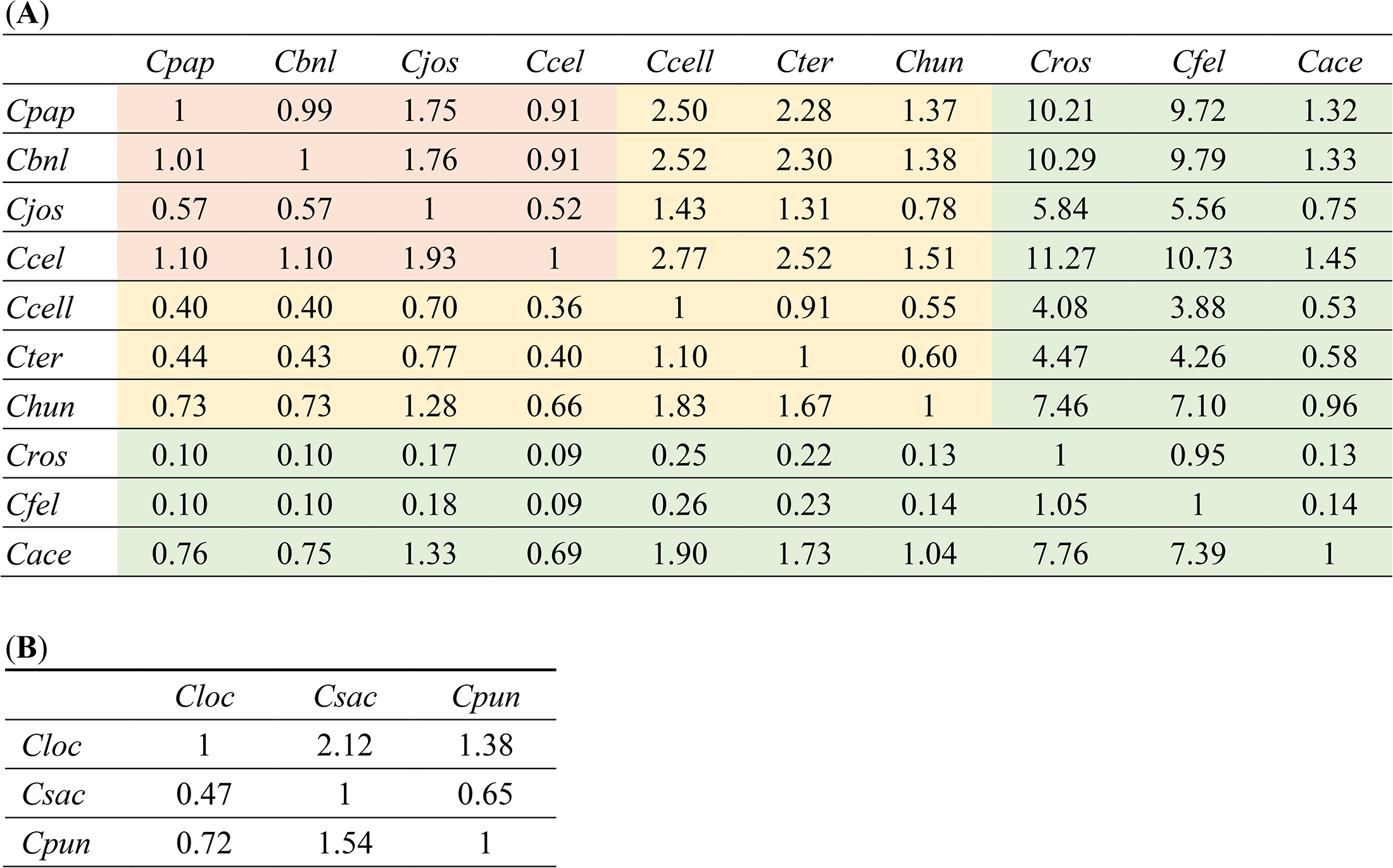
Evolutionary ratio (ER) matrix for the cellulosome complex operons from the 13 Clostridial species. (**A**) ERs for Clade-1 of CoSL-tree, where the subclades of *Cpap-Cbnl-Cjos-Ccel*, *Ccell-Cter-Chun* and *Cros-Cfel-Cace* are colored with orange, yellow and green respectively. (**B**) ERs for Clade-2 of CoSL-tree.

In Subclade 1.1, the SLs in *Ccell* operon are more similar to *Cter* than to *Chun*, *i.e.*, the operon ERs for *Cter-Ccell* and *Chun-Ccell* are 1.10 and 1.83 respectively (**Table 1A**), while those for *Cter-Chun* and *Ccell-Chun* are 0.60 and 0.55 respectively. Thus *Ccell* and *Cter* are in purifying selection, while Subclade 1.1 is under positive selection towards *Chun* (similar to as Subclade 1.2; **Fig. 1C**; **Fig. S5**).

The Clade 2 species in CoSL-tree are more dynamic in evolution than Clade 1, in that they show more variable number of genes and SLs. *Cloc*, *Csac* and the out-grouped *Cpun* exhibit a certain degree of similarity to the Clade 1 species, but feature the addition of new SLs such as SL-3A and SL-7A (homologous to SL3 and SL7 respectively; **Fig. 1C**). Moreover, their cellulosomal operons are distinct, e.g., *Cpun* and *Csac* operons harbor no cohesin, glycoside hydrolase (GH) or dockerin genes. In fact, ERs for *Cpun* and *Csac* versus *Cloc* are 1.37 and 2.11 respectively (**Table 1B**), consistent with positive selection.

Notably, if the ERs are calculated without considering SLs (and the CV) in **Equation I**, then the number of genes by itself is not sufficient to detect the selection. For example, in Subclade 1.2 (*Cpap-Cbnl-Cjos-Ccel*; *Cros-Cfel-Cace*), the equal number of genes would suggest an ER of 1, which however is misleading. Therefore, in computing CoSL-based ER, the SLs are essential for deriving ERs in CoSLOE.

### 3.5 The CoSLOE model of cellulosomal operons is supported by variation in enzyme genes

In CoSLOE, purifying/negative selection occurs when the tree is traversed from the root to the leaf nodes, and diversifying/positive selection takes place when traversing from leafs to the root nodes (**Fig. 4**). In the cellulosomal operon (**Fig. 1C**), positive evolution takes place in the *Ccel-Cjos-Cbnl-Cpap* direction (root to leaf), in the *Chun-Cter-Ccell* direction and in the *Cace-Cros-Cfel* direction respectively, with the root node being the most positively selected and the leaf nodes the most negatively selected.

**Figure 4.**
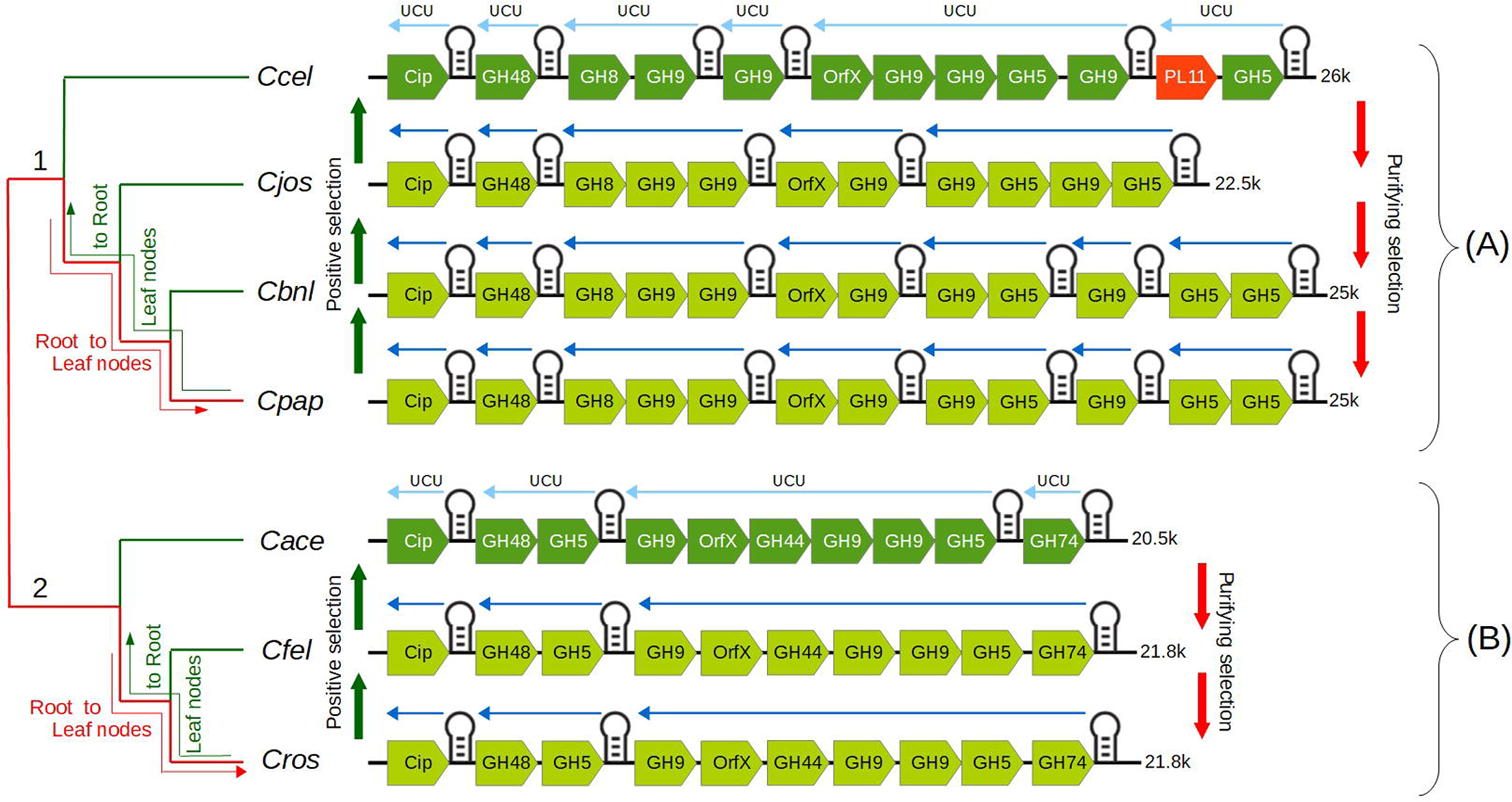
Evolution of the cellulosome-encoding SRPS operons based on CoSLOE. The two clades in the tree represent two different operon evolution scenarios yet with an identical evolutionary directional flow, *i.e.*, the movement from root to leaf nodes defines purifying (i.e., negative) selection and the reverse movement (i.e., leaf to root) depicts diversifying selection (i.e., positive selection). Upstream Controlled Unit (UCU) represents the region (which is upstream to a SL and can include multiple genes) that is regulated by a SL via the SRPS mechanism. (**A**) In Clade 1 (with four species), positive selection resulted in distinct ΔG of SLs (each corresponding to a UCU) in the outermost operon of the clade, while those operons with similar ΔG are conserved in the leaf nodes. Variation in a UCU can be caused by gain/loss of the SLs along with the corresponding genes (i.e., depicting appearance and disappearance of new genes). (**B**) In Clade 2 (with three species), positive selection also led to distinct ΔG of SLs (each corresponding to a UCU) in the outermost species, suggesting identical evolutionary flow in both clades. However, the positively selected operons carry characteristics distinct from the leaf-node operons, despite an identical number of SLs and genes. Thus the change in the ΔG of SLs can lead to operons with discrete function. Color gradient of genes represents positive (darker color) or purifying (light color) selection.

To probe the biological significance of these findings, the genome-wide numbers of carbohydrate-active enzymes (CAZymes) and carbohydrate-binding module (CBM) were compared, since these enzymes are major parts of the cellulosomal system(Busch *et al.*, 2017). For example, in Subclade 1.2, for the *Ccel*, *Cjos*, *Cbnl* and *Cpap* genome (which exhibit > 95% similarity in 16S rRNA sequences; **Fig. 1C, 6A**), (*i*) the CAZymes (including glycoside hydrolases or GHs, carbohydrate esterases or CEs, and polysaccharide lyases or PLs) harbored is 111 (94 GHs, 13 CEs, 4 PLs), 116 (92 GHs, 19 CEs, 5 PLs), 127 (103 GHs, 19 CEs, 5 PLs) and 122 (103 GHs, 16 CEs, 3 PLs) respectively (Dassa *et al.*, 2017), exhibiting an overall pattern of increase; (*ii*) for GH5 (*Ccel*: 7; *Cjos*: 7; *Cbnl*: 8; *Cpap*: 7), GH9 (13, 14, 14, 14) and GH43 (9, 13, 13, 13), an increase in number is apparent when traversing from root to the leave nodes (*Ccel-Cjos-Cbnl-Cpap*); (*iii*) a similar pattern (i.e., increase in number) is observed in CBMs (54, 59, 67, 71) and to a less degree, dockerins (69, 72, 88, 68) (Dassa *et al.*, 2017) (**Fig. 4A**). Thus *Ccel* is an outlier in terms of the genome-wide CAZyme number. Moreover, of 26Kb in size, the cellulosomal operon of *Ccel* is the largest (**Fig. S5**; *Cjos*: 22.5Kb; *Cbnl*: 25Kb; *Cpap*: 25Kb), and harbors unique genes such as pectin degrading enzymes (Pagès *et al.*, 2003, McDonald *et al.*, 2008) (*Rgl11Y*) and longer hybrid linkers (Pinheiro *et al.*, 2008) that join cohesins to scafoldins. Similarly, in the *Cace-Cfel-Cros* cluster, the *Cace* cellulosomal operon harbors one additional enzyme (Sialidase; **Fig. 4B**) yet lacks cellulosomal complex activity (Sabathé *et al.*, 2002), in opposite to *Cros* and *Cfel* which are used for the retting process (Angelini *et al.*, 2013). These observations are consistent with CoSLOE-derived positive selection of the *Ccel* cellulosomal operon.

Similarly, in Subclade 1.1, the cellulosomal operon of *Chun* uniquely harbors a xyloglucanse gene. Thus the near-root cellulosome operons are positively selected towards less efficient cellulosic activity or addition of auxiliary functionality, supporting CoSLOE-derived operon selection.

### 3.6 Evolution of the ATP synthase operons via CoSLOE is similar to the cellulosomal operons

CoSL-tree of the ATP synthase operons is similar to that of the cellulosome operons (**Fig. 5A**; **Fig. 1C**), except that *Cpun* is clustered with *Chun* in the former. Notably, within each of the *Chun-Cell-Cter*, *Ccel-Cpap-Cbnl-Cjos* and *Cros-Cace-Cfel* subclades, operon sequences are nearly 100% similar, and the gene sequences of subunit alpha and beta are conserved across 13 species (**Fig. 5A**). The less variation in gene sequences (than cellulosome operon) among 13 *Clostridium* species is probably due to the strict functional conservation of the ATP synthase complex.

**Figure 5.**
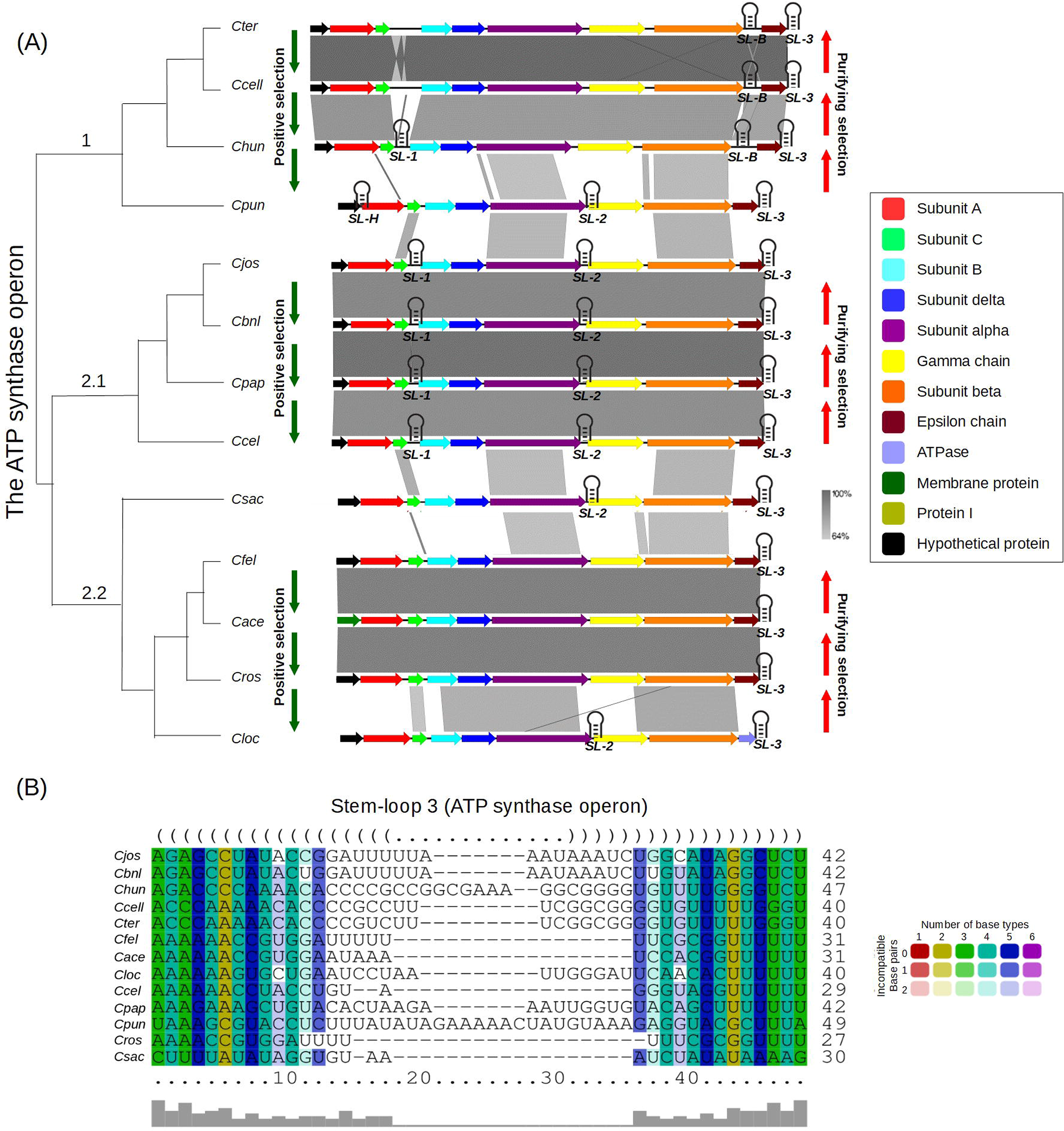
Evolution of the ATP-synthase-encoding SRPS operons in the 13 Clostridial species based on the CoSLOE model. (**A**) ATP synthase operons sequences were mapped sequence-wise according to the CoSL-based phylogeny, where the sequence similarity is shown by dark black (100% similarity) and light black (64% similarity) color gradient and the genes are colored based on their encoding protein. The ERs determine the positive and negative selection pattern (shown via green and red arrows, respectively). (**B**) The multiple sequence alignment of SL-3 from the ATP synthase operons in the 13 Clostridial species.

As in cellulosome operons, gene-SL relationship was probed in the ATPase operons. Three orthologous SLs were predicted in ATPase operon, where (*i*) SL-1 is preserved in *Chun, Cjos, Cbnl, Cpap* and *Ccel* (always flanking at 3’ UTR of subunit C); (*ii*) SL-2 is present at 3’ UTR of subunit alpha in *Cpun, Cjos, Cbnl, Cpap, Ccel,* and *Csac*; (*iii*) SL-3 is conserved throughout the 13 species at the 3’ UTR of epsilon chain and terminating the operon; (*iv*) Only one SL is predicted in *Cace, Cfel, Cros*, and *Cloc*, suggesting that their ATP synthase operons seem not regulated by the SRPS mechanism (**Fig. 5B**). Taken together, these associations between genes and SLs show their relationship, which is consistent with the observation in cellulosome operons.

As for ER, in Clade 1, the ERs for *Cter-Cpun*, *Ccell-Cpun* and *Chun-Cpun* are 0.34, 0.41 and 0.80 respectively (**Table S3**), revealing negative selection towards leaf nodes in Clade 1 (**Fig. 5A**), *i.e. Ccell* and *Cter* appear to undergo purifying selection, while *Chun* is positively selected towards *Cpun* (ER for *Cpun-Chun*: 1.24; **Table S3**). In Subclade 2.1, similarly, ERs of *Cjos-Ccel*, *Cbnl-Ccel* and *Cpap-Ccel* are 0.94, 0.93, 1.20, respectively (**Table S3**), i.e. the overall flow of *Cjos-Cbnl-Ccel* is consistent with positive selection, except for *Cpap* (**Fig. 5A**).

Interestingly, the ATP synthase operons exhibit an evolution pattern similar to the cellulosome operons, by positive selection in the *Cter-Ccell-Chun* and *Cjos-Cbnl-Cpap-Ccel,* direction (purifying selection in the reverse direction; **Fig. 5A**). Since ATP synthase operon is functionally conserved in most species (Neupane *et al.*, 2019), less variability was present in the genes and SLs. However, the root species of Subclade 1.1 (*Chun*), 1.2 (*Ccel*) and 2.1 (*Cloc*) harbors smaller operons, longer operons and an additional enzyme at the 3’ UTR region, respectively (**Fig. 5A**), consistent with the evolutionary pattern suggested by the observations in cellulosome operons.

### 3.7 Genome-wide application of CoSLOE reveals the direction of organismal selection

The evolutionary flow in a tree represents the different directions that species follow due to the selection-pressure on them, during evolution. To probe SL-driven evolutionary selection-pressure, orthologous SRPS operons were probed using CoSLOE. However, due to the lack of orthology among SRPS operons in 13 species, operon evolution was probed clade-wise in CoSL-tree (**Fig. 1C**), *i.e.* Subclade 1.2 (*Ccel-Cjos-Cbnl-Cpap*). In Subclade 1.2, orthologous SRPS operons are scattered across the genomes of *Ccel*, *Cjos*, *Cbnl* and *Cpap* which are in the form of chromosome *(Cbnl* and *Ccel*) or contigs (*Cpap*-31 and *Cjos*-2; **Fig. 6A**; **Table S4**).

**Figure 6.**
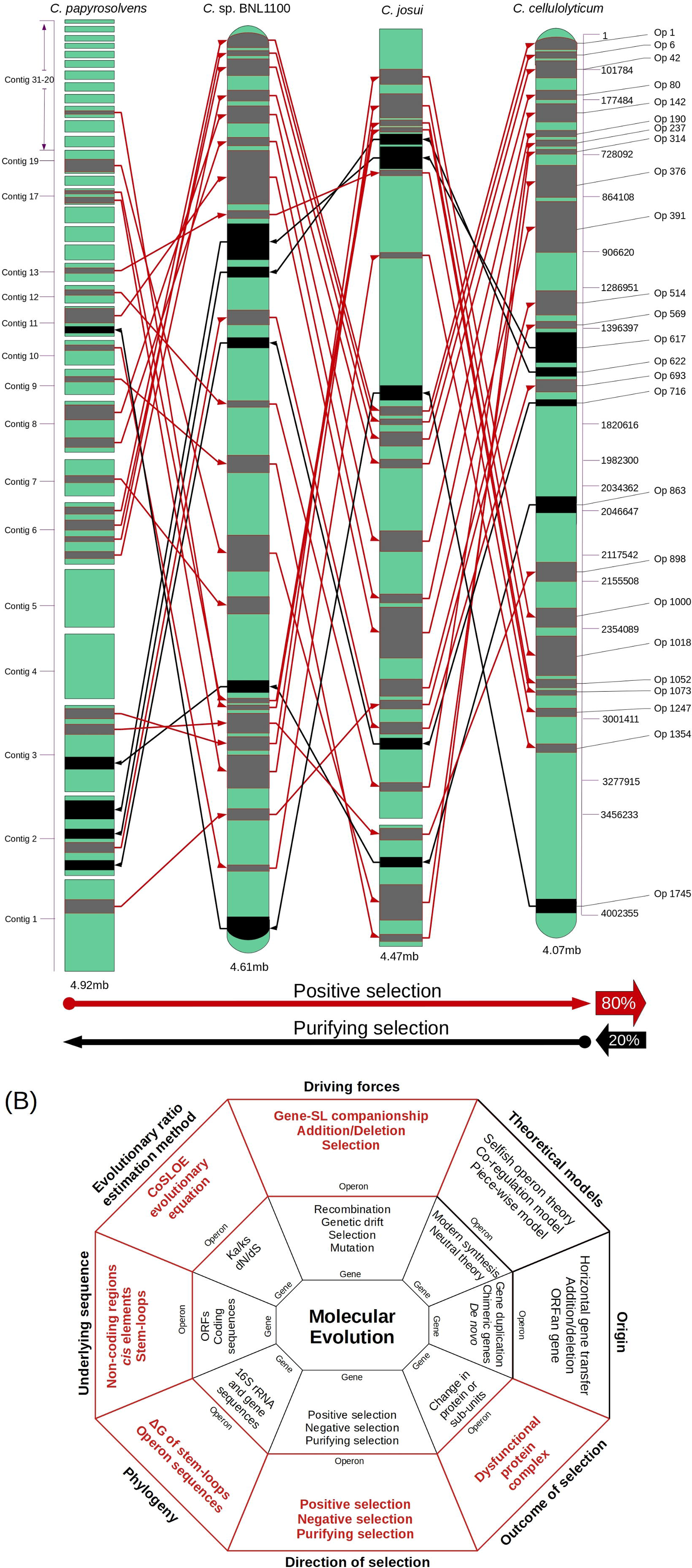
The genome-wide CoSLOE model defines the direction of organismal selection. (**A**) Genome-wide evolution of SRPS operons based on the CoSLOE model (**Table S4**). Orthologous SRPS operons are scattered across the genomes of *C. cellulolyticum* (*Ccel*), *C. josui* (*Cjos*), *C.* sp. BNL1100 (*Cbnl*), and *C. papyrosolvens* (*Cpap*), represented here in the form of chromosome (*Cbnl* and *Ccel*) and contigs (*Cpap*-31 and *Cjos*-2). Totally, five out of 25 SRPS operons follow the *Ccel-Cjos-Cbnl-Cpap* direction and the other 80% SRPS operons show a positive selection flow in the *Cpap-Cbnl-Cjos-Ccel* direction, *i.e*., from the leaf nodes to the root nodes. Dark and grey region/band represents operon (Op), and their thickness shows the length. Direction of arrows represents direction of selection pressure, either positive (red) or negative (black). (**B**) Link and distinction between CoSLOE and the existing models for operon evolution. Information derived from the CoSLOE model is highlighted in red.

Here, five out of the 25 SRPS operons, *i.e.*, Op617, Op622, Op716, Op863 and Op1745, follow the *Ccel-Cjos-Cbnl-Cpap* direction (black arrows; **Fig. 6A**). Interestingly, the other 80% SRPS operons show the positive selection flow in the *Cpap-Cbnl-Cjos-Ccel* direction, *i.e.* from the leaf nodes to the root nodes, which is similar to the cellulosomal cluster evolutionary flow, e.g., for Op142, Op376 and Op898 (red arrows; **Fig. 6A**). These observations suggest that the SRPS mechanism, although evolutionarily conserved, can reveal selection-pressure that is distinct from organismal phylogeny.

## 4 Discussion and conclusion

In existing frameworks of operon evolution, coding sequences (i.e., subunits of protein complex or components of metabolic pathway encoded by the operon) have been thought to play a major role. They can drive structural variation and functional adaptation of operons towards a specific niche (Gogarten *et al.*, 2002, Francino *et al.*, 2012), for example, by deletion or insertion of the whole genes or via synonymous/non-synonymous mutations of their sequences. However, it remains elusive whether non-coding elements play a role in such adaptation of operons.

The stoichiometry of SRPS operons, found genome-wide, can be modeled based on the genome sequence of SLs alone (Bhaskar *et al.*, 2021), suggesting a quantitative model of evolution at the whole operon level, in parallel to the evolution at the individual coding gene level (e.g., Ka/Ks (Kimura *et al.*, 1968)). Based on ΔG of such SLs, we proposed CoSLOE, with evolutionary ratio (ER) >1 or <1 for positive or negative selection of SRPS operons. In the CoSL-tree of cellulosomal operons, when traversing from leafs to the root nodes, ERs reveal diversifying/positive selection towards a less efficient cellulosomal system, consistent with glycoside-hydrolase gene variation both in-operon and genome-wide. A consistent pattern is followed by the ATPase operon and the majority of orthologous SRPS operons genome-wide, suggesting conservation among operons in such selection. Therefore, CoSLOE provides a new layer of insights into operon evolution that is distinct from existing models (**Fig. 6B**). Specifically, (*i*) Driving forces: for individual genes, mutation, recombination, genetic drift and selection are known evolution drivers; for SRPS operons, in addition to addition/deletion/mutation of genes, CoSLOE introduces SLs a previously unrecognized driver. (*ii*) Theoretical models: in addition to the known models of gene evolution (neutral theory (Nei *et al.*, 2005)) and operon evolution (selfish operon theory (Lawrence & Roth *et al.*, 1996), co-regulation model (Price *et al.*, 2005) and piece-wise model (Fani *et al.*, 2005)), CoSLOE provides a new framework for quantitatively modeling SRPS operon evolution. (*iii*) Rate of selection: CoSLOE compares rate of selection between two orthologous operons, which is conceptually similar to Ka/Ks or dN/dS which compares between orthologous genes. (*iv*) Outcome of selection: gene evolution generally results in changed protein sequence, yet the operon evolution depicted by CoSLOE results in altered ingredient or stoichiometry of the whole protein complex or metabolic pathway. (*v*) Direction of selection: just like Ka/Ks for orthologous genes, CoSLOE offers strategy to model the direction for orthologous operons. (*vi*) Phylogeny: a tree based on ΔG of SLs of SRPS operons can model the selection of operon and organism, in contrast to 16S-rRNA gene trees that model organismal phylogeny. (*vii*) Underlying sequence: instead of relying on coding sequences, CoSLOE takes advantage of the non-coding sequences to model operon evolution, and highlights the role of *cis*-elements in shaping operon evolution and organism adaptation. (*viii*) Origin of operons: CoSLOE suggests the SLs (and their relationship with associated genes) as a key player in the original formation of operon structure, in addition to horizontal gene transfer, deletion of intervening genes and addition of ORFan genes (Price *et al.*, 2006).

Notably, we have tested CoSLOE on just 13 Clostridial species, and expansion of the model to a broader range of species is limited by the paucity of experimental data and lack of computational approaches to identify SPRS operons. Therefore, to what degree the model is applicable across microorganisms is not yet clear, and answer to this question is perhaps ultimately dependent on the breath and boundary of SPRS mechanism. Despite these limitations, for SRPS operons, our findings here reveal the link between operon stoichiometry and operon evolution, and propose a new *cis*-element-based framework to model the direction and rate of SRPS operon evolution.

## 5 Acknowledgements

This work was supported by University of Chinese Academy of Sciences Scholarship for International PhD students.

## 6 Author contribution

YB and JX designed the study; YB performed the computational analysis; YB and JX analyzed the data; MHB and CX provided critical suggestions; YB and JX wrote the paper.

## 8 Data availability

The data underlying this article are available in the article and in its online supplementary material.

## 12 Supplementary Tables and Figures

**Table S1. Free energy (ΔG) of harbored SLs in the SRPS operons that encode cellulosome from 13 Clostridial species.**

**Table S2. Ka/Ks values for the first gene in the cellulosomal operon from *Cpap*,*Cbnl*,*Cjos* and *Ccel*.**

**Table S3. The evolutionary ratio (ER) matrix for the ATP synthase operons from the 13 Clostridial species.**

**Table S4. The number of genes and SLs in all SRPS operons in the *Ccel*, *Cjos*, *Cbnl* and *Cpap* genomes.**

**Figure S1. Phylogenetic tree of 13 Clostrdial species based on the predicted ΔG of the SLs (CoSL-tree) (A) or the 16S rRNA sequences (16S-tree) (B).**

**Figure S2. Clade-wise representation of the SL-1 (A), SL-2 (B), SL-3 (C), SL-4 (D), SL-5 (E), SL-6 (F), SL-7 (G), SL-3A (H), SL-2A (I) and SL-7A (J), based on their sequence and structure similarity.**

**Figure S3. Linear comparison of the nucleic acid sequences of the cellulosome operons from the 13 Clostridial species.**

