## Supplementary Tables and Figures for "Evolution of Selective RNA Processing and Stabilization operons in cellulosome-harboring *Clostridium* spp"

This file contains the following materials:

1. Supplementary figures.
2. Supplementary tables.

### Supplementary figures

**(A)**

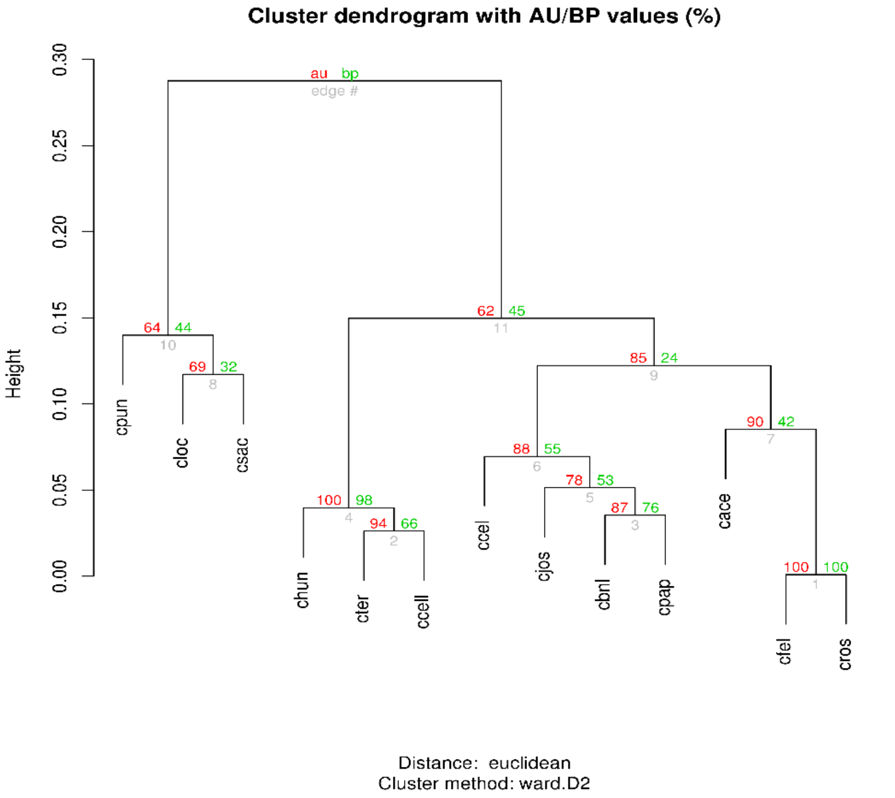

**(B)**

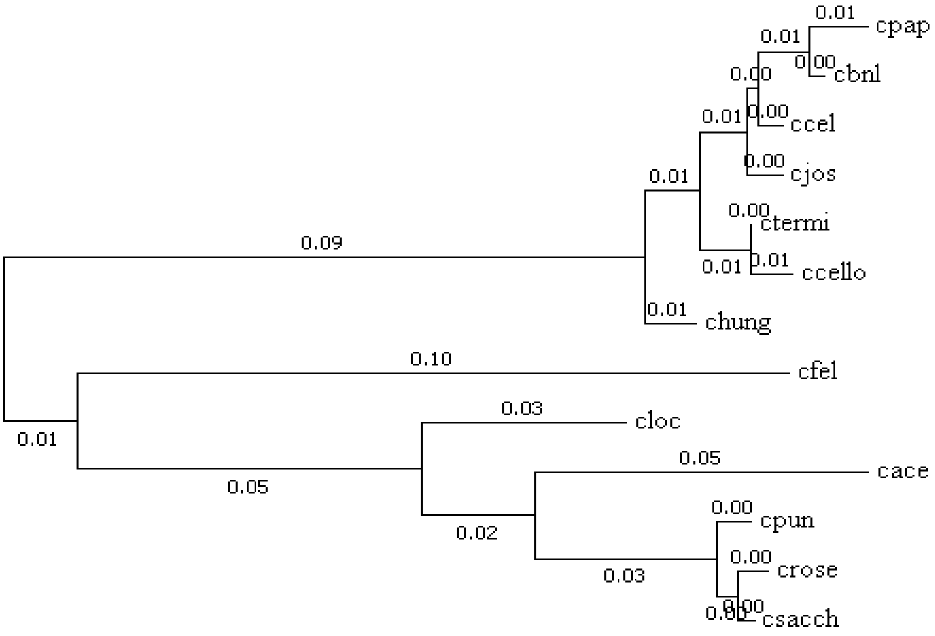

**Figure S1. Phylogenetic tree of 13 Clostrdial species based on the predicted ΔG of the SLs (CoSL-tree) (A) or the 16S rRNA sequences (16S-tree) (B).**

**(A)** Stem-loop 1

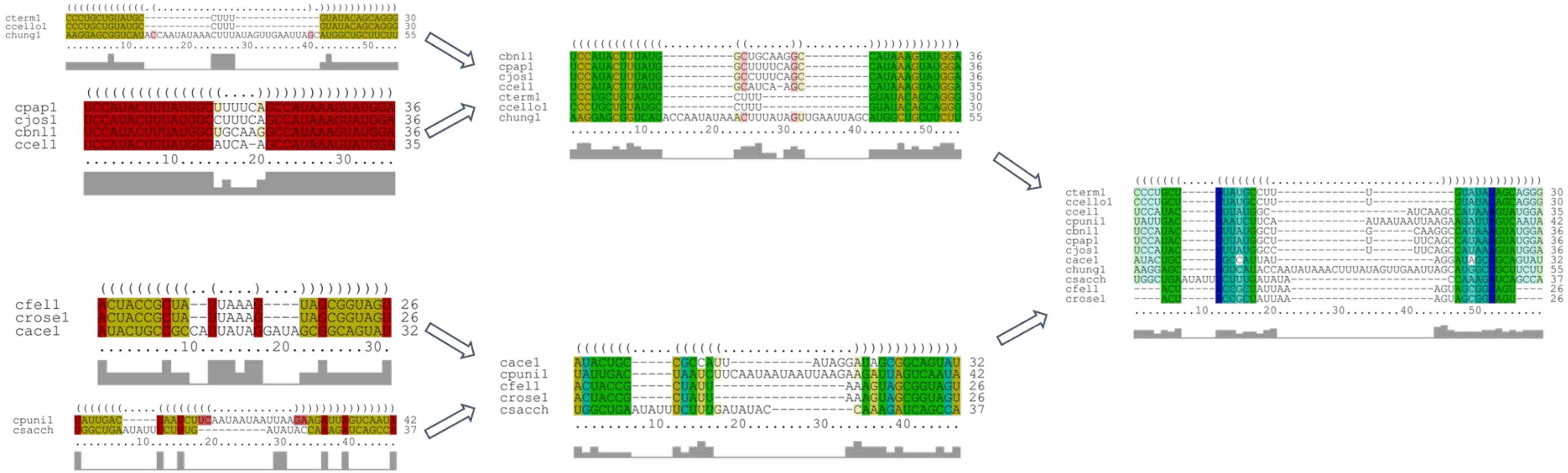

**(B)** Stem-loop 2

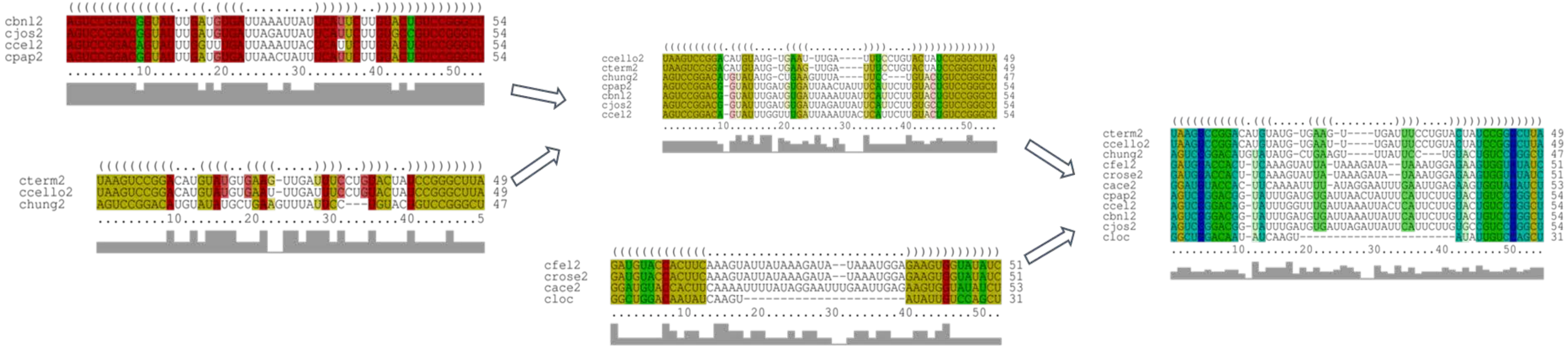

(**C**) Stem-loop 3

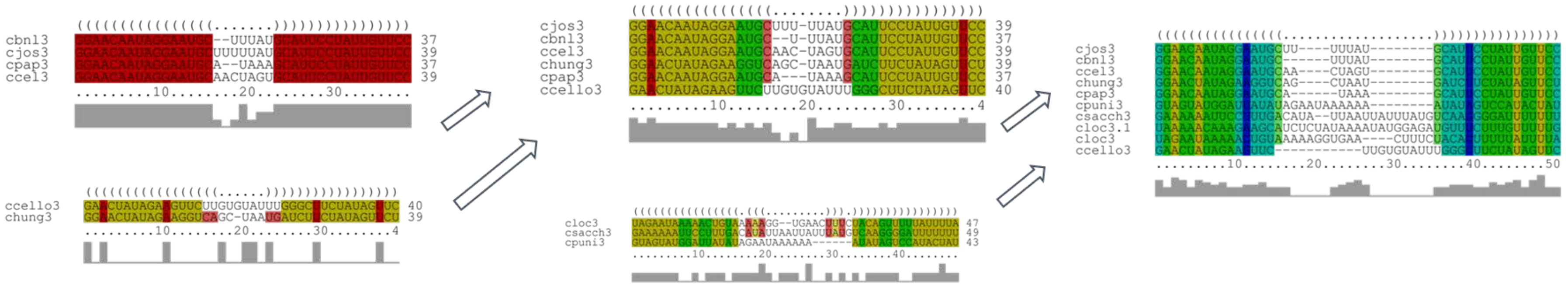

(**D**) Stem-loop 4

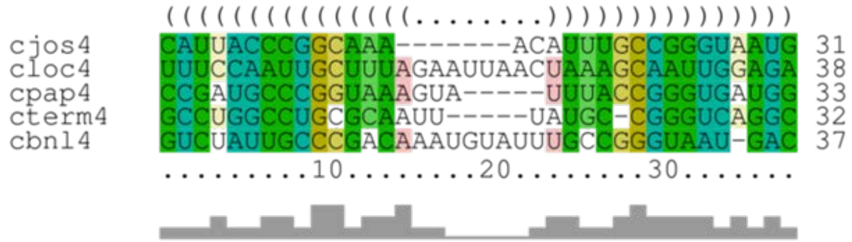

(**E**) Stem-loop 5

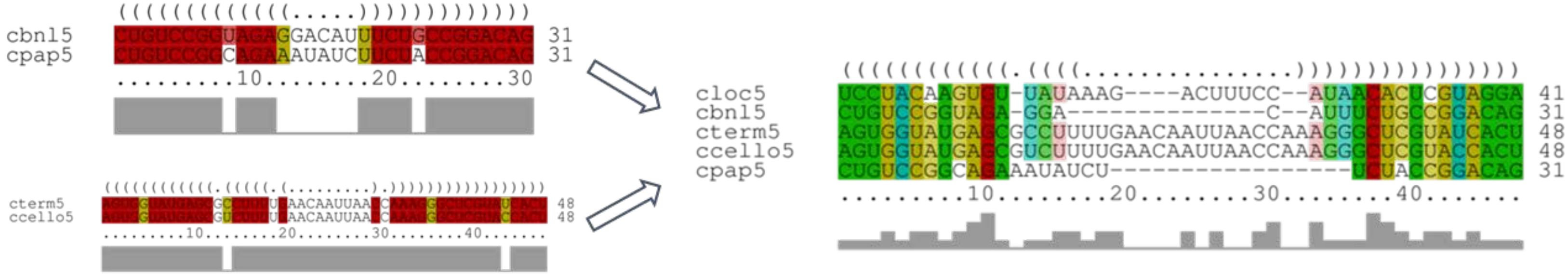

**(F)** Stem-loop 6

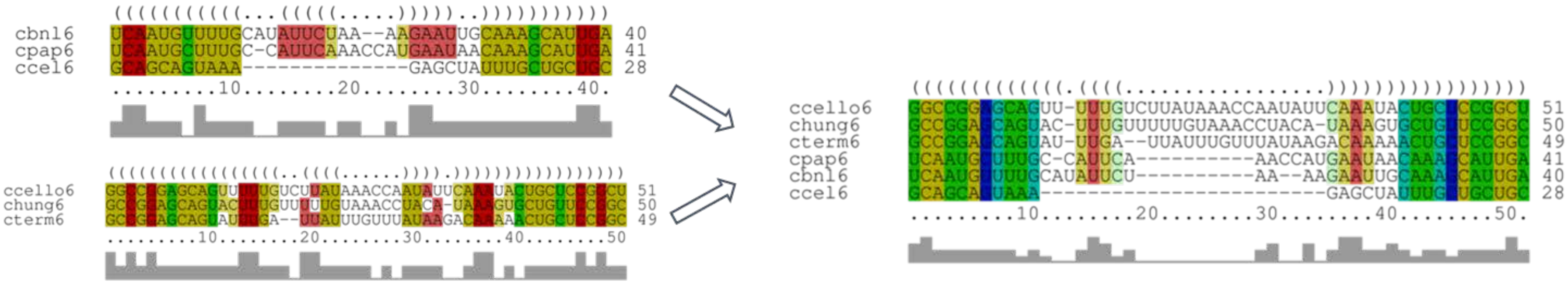

**(G)** Stem-loop 7

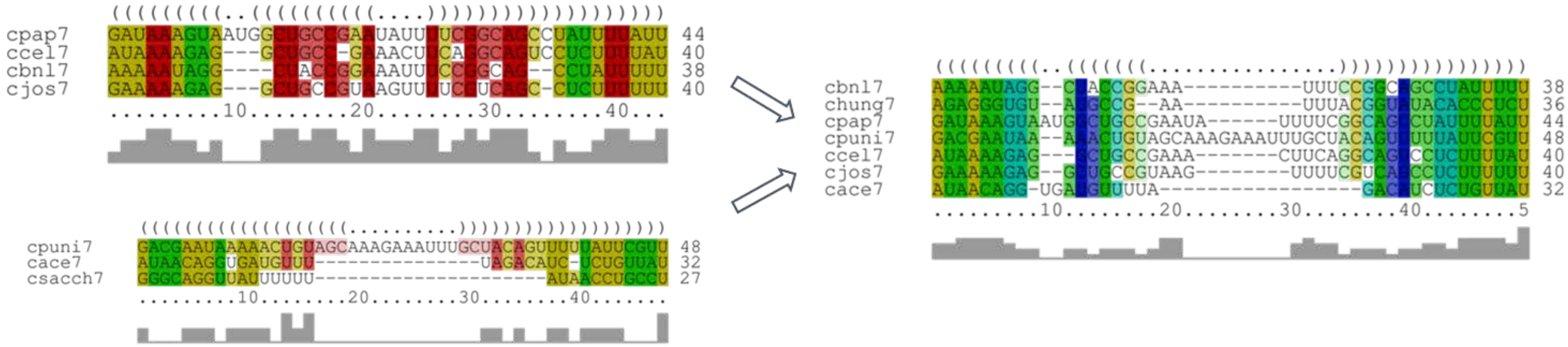

(**J**) Stem-loop 3A

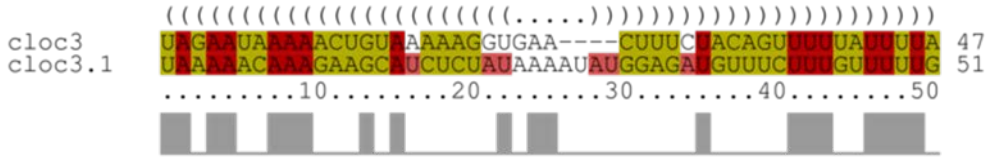

(**I**) Stem-loop 2A

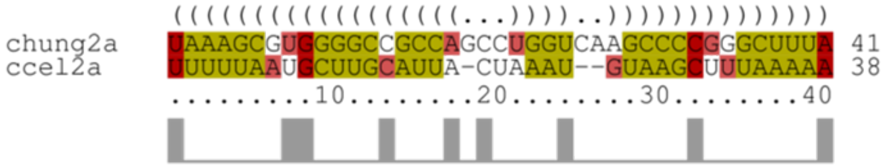

(**H**) Stem-loop 7A

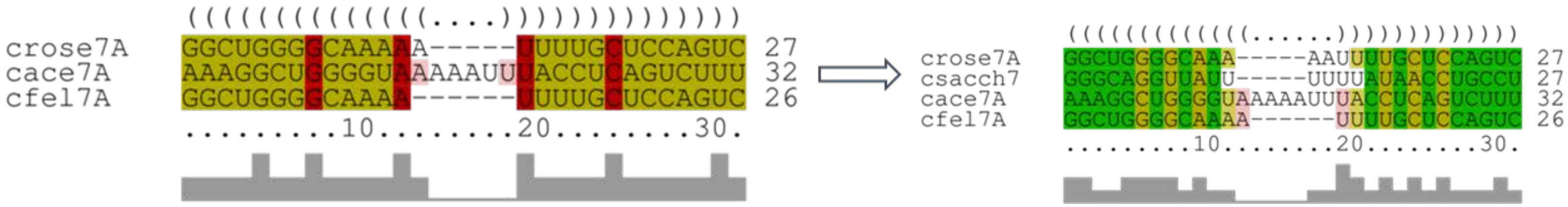

**Figure S2. Clade-wise representation of the SL-1 (A), SL-2 (B), SL-3 (C), SL-4 (D), SL-5 (E), SL-6 (F), SL-7 (G), SL-3A (H), SL-2A (I) and SL-7A (J), based on their sequence and structure similarity.**

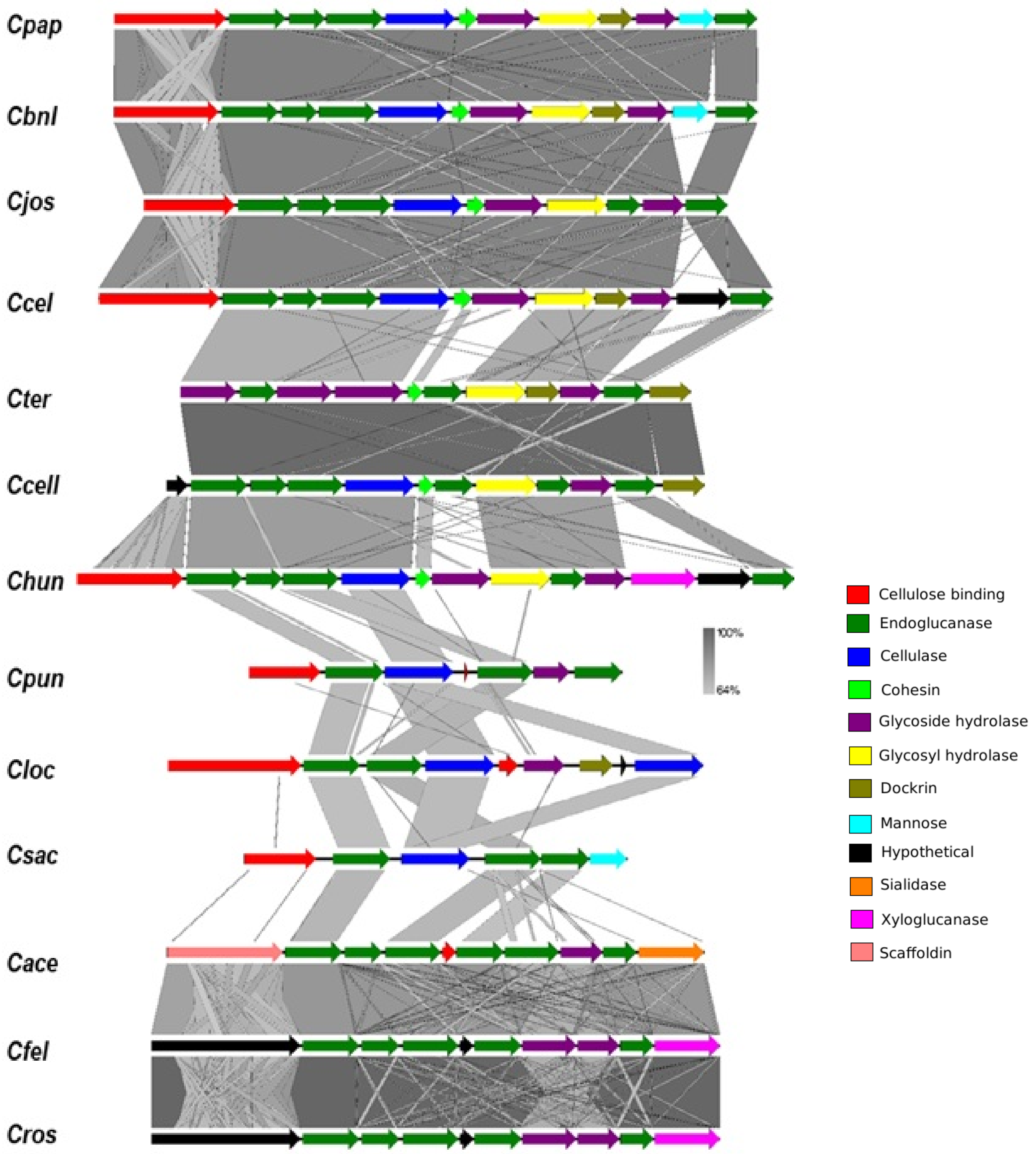

**Figure S3. Linear comparison of the nucleic acid sequences of the cellulosome operons from the 13 Clostridial species.**

### Supplementary Tables:

**Table S1. Free energy (ΔG) of harbored SLs in the SRPS operons that encode cellulosome from 13 Clostridial species.**

| Organism | Number of genes | Stem-loops predicted | ΔG-based ratio | Normalized ratio |
|---|---|---|---|---|
| *Clostridium* sp. BNL1100 (*Cbnl*) | 12 | 7 | -24.4: -26.3: -25.9: -25.9: -25.9: -15.3: -15.3: -21.2: -21.2: -18.3: -21.5: -21.5 | 0.0929: 0.1001: 0.0986: 0.0986: 0.0986: 0.0582: 0.0582: 0.0807: 0.0807: 0.0697: 0.0818: 0.0818 |
| [*Clostridium*] *papyrosolvens* (*Cpap*) | 12 | 7 | -23.6: -26.3: -25.3: -25.3: -25.3: -16.7: -16.7: -16.8: -16.8: -23.5: -23.9: -23.9 | 0.0892: 0.0994: 0.0957: 0.0957: 0.0957: 0.0631: 0.0631: 0.0635: 0.0635: 0.0904: 0.0904: 0.0904 |
| [*Clostridium*] *josui* (*Cjous*) | 11 | 5 | -22.6: -29.5: -25.6: -25.6: -25.6: -21.3: -21.3: -19.1: -19.1: -19.1: -19.1 | 0.0912: 0.1190: 0.1033: 0.1033: 0.1033: 0.0859: 0.0859: 0.0770: 0.0770: 0.0770: 0.0770 |
| [*Clostridium*] *cellulolyticum* (*Ccel*) | 12 | 6 | -23.5: -26.8: -14.5: -14.5: -26.2: -16.3: -16.3: -16.3: -16.3: -16.3: -20.9: -20.9 | 0.1027: 0.1171: 0.0634: 0.0634: 0.1145: 0.0712: 0.0712: 0.0712: 0.0712: 0.0712: 0.0913: 0.0913 |
| [*Clostridium*] *termitidis* (*Cter*) | 11 | 4 | -23.1: -18: -21.1: -21.1: -25.6: -25.6: -25.6: -25.6: -25.6: -25.6: -21.8: -21.8 | 0.0824: 0.0642: 0.0752: 0.0752: 0.0913: 0.0913: 0.0913: 0.0913: 0.0913: 0.0913: 0.0777: 0.0777 |
| [*Clostridium*] *cellobioparum* (*Ccell*) | 12 | 5 | -21.7: -18.7: -18.9: -18.9: -18.9: -23.5: -23.5: -23.5: -23.5: -23.5: -23.2: -23.2 | 0.0831: 0.0716: 0.0724: 0.0724: 0.0724: 0.0900: 0.0900: 0.0900: 0.0900: 0.0900: 0.0889: 0.0889 |

| | | | | |
|---|---|---|---|---|
| [*Clostridium*] *hungatei* (*Chun*) | 13 | 6 | -18: -20.8: -23.2: -23.2: -23.3: -28.6: -28.6: -28.6: -28.6: -28.6: -23.3: -23.3: -23.3 | 0.0560: 0.0647: 0.0722: 0.0722: 0.0725: 0.0890: 0.0890: 0.0890: 0.0890: 0.0890: 0.0725: 0.0725: 0.0725 |
| *Clostridium cellulovorans* (*Cloc*) | 9 | 5 | -18.2: -18.2: -18.3: -18.3: -21.2: -21.2: -15.2: -28.4: -28.4 | 0.0971: 0.0971: 0.0977: 0.0977: 0.1131: 0.1131: 0.0811: 0.1515: 0.1515 |
| *Clostridium saccharoperbutylacetonicum* (*Csac*) | 8 | 3 | -14.4: -23.1: -23.1: -23.1: -16.5: -16.5: -16.5: -16.5 | 0.0962: 0.1543: 0.1543: 0.1543: 0.1102: 0.1102: 0.1102: 0.1102 |
| *Clostridium puniceum* (*Cpun*) | 7 | 3 | -17.5: -17.4: -17.4: -22.7: -29.6: -29.6: -29.6 | 0.1003: 0.0998: 0.0998: 0.1302: 0.1697: 0.1697: 0.1697 |
| *Clostridium acetobutylicum* (*Cace*) | 10 | 4 | -17: -22.6: -22.6: -12.2: -12.2: -12.2: -12.2: -12.2: -12.2: -18.6 | 0.1104: 0.1468: 0.1468: 0.0792: 0.0792: 0.0792: 0.0792: 0.0792: 0.0792: 0.1208 |
| *Clostridium felsineum* (*Cfel*) | 10 | 3 | -16.6: -18.5: -18.5: -16.4: -16.4: -16.4: -16.4: -16.4: -16.4: -16.4 | 0.0986: 0.1099: 0.1099: 0.0974: 0.0974: 0.0974: 0.0974: 0.0974: 0.0974: 0.0974 |
| *Clostridium roseum* (*Cros*) | 10 | 3 | -16.6: -18.5: -18.5: -16.5: -16.5: -16.5: -16.5: -16.5: -16.5: -16.5 | 0.0982: 0.1094: 0.1094: 0.0976: 0.0976: 0.0976: 0.0976: 0.0976: 0.0976: 0.0976 |

**Table S2. Ka/Ks values for the first gene in the cellulosomal operon from *Cpap*, *Cbnl*, *Cjos* and *Ccel*.**

| | | | | |
|---|---|---|---|---|
| *Cpap*1 | 1 | | | |
| *Cjos*1 | 1.508 | 1 | | |
| *Cbnl*1 | 0.543 | 1.453 | 1 | |
| *Ccel*1 | 0.589 | 1.424 | 0.329 | 1 |

**Table S3. The evolutionary ratio (ER) matrix for the ATP synthase operons from the 13 Clostridial species.**

| | *Cpap* | *Cbnl* | *Cjos* | *Ccel* | *Ccell* | *Cter* | *Chun* | *Cpun* |
|---|---|---|---|---|---|---|---|---|
| *Cpap* | 1 | 1.288 | 1.270 | 1.202 | 1.723 | 2.081 | 0.893 | 0.717 |
| *Cbnl* | 0.776 | 1 | 0.986 | 0.933 | 1.338 | 1.615 | 0.693 | 0.557 |
| *Cjos* | 0.787 | 1.014 | 1 | 0.947 | 1.357 | 1.639 | 0.703 | 0.565 |
| *Ccel* | 0.831 | 1.071 | 1.056 | 1 | 1.433 | 1.731 | 0.743 | 0.597 |
| *Ccell* | 0.580 | 0.748 | 0.737 | 0.698 | 1 | 1.208 | 0.518 | 0.416 |
| *Cter* | 0.480 | 0.619 | 0.610 | 0.578 | 0.828 | 1 | 0.429 | 0.345 |
| *Chun* | 1.120 | 1.443 | 1.422 | 1.347 | 1.930 | 2.330 | 1 | 0.803 |
| *Cpun* | 1.394 | 1.796 | 1.770 | 1.676 | 2.402 | 2.901 | 1.245 | 1 |

**Table S4. The number of genes and SLs in all SRPS operons in the *Ccel*, *Cjos*, *Cbnl* and *Cpap* genomes.** Operon number is shown for *C. cellulolyticum*.

| Operon | *C.cellulolyticum* | | *C. josui* | | *C.* sp. BNL1100 | | *C. papyrosolvens* | |
|---|---|---|---|---|---|---|---|---|
| | Genes | Stem-loops | Genes | Contig | Genes | Stem-loops | Genes | Contig |
| 1 | 4 | 1 | 4 | NZ_JAGE01000001.1 | 4 | 2 | 4 | NZ_ACXX02000006.1 |
| 6 | 3 | 2 | 3 | NZ_JAGE01000001.1 | 4 | 1 | 3 | NZ_ACXX02000006.1 |
| 42 | 9 | 1 | 7 | NZ_JAGE01000001.1 | 8 | 1 | 8 | NZ_ACXX02000006.1 |
| 80 | 4 | 2 | 4 | NZ_JAGE01000001.1 | 4 | 0 | 4 | NZ_ACXX02000006.1 |
| 142 | 8 | 3 | 8 | NZ_JAGE01000001.1 | 8 | 3 | 8 | NZ_ACXX02000008.1 |
| 190 | 4 | 2 | 4 | NZ_JAGE01000001.1 | 4 | 1 | 4 | NZ_ACXX02000008.1 |
| 216 | 3 | 1 | - | - | - | - | - | - |
| 237 | 3 | 1 | 3 | NZ_JAGE01000001.1 | 2 | 1 | 2 | NZ_ACXX02000012.1 |
| 288 | 5 | 1 | - | - | - | - | - | - |
| 314 | 3 | 2 | 3 | NZ_JAGE01000002.1 | 3 | 2 | 2 | NZ_ACXX02000009.1 |
| 376 | 12 | 6 | 11 | NZ_JAGE01000002.1 | 12 | 7 | 12 | NZ_ACXX02000019.1 |
| 391 | 24 | 2 | 24 | NZ_JAGE01000001.1 | 24 | 2 | 24 | NZ_ACXX02000011.1 |
| 495 | 3 | 2 | - | - | - | - | - | - |
| 511 | 9 | 1 | - | - | - | - | - | - |
| 514 | 11 | 2 | 8 | NZ_JAGE01000001.1 | - | - | - | - |
| 569 | 3 | 1 | 3 | NZ_JAGE01000001.1 | 3 | 1 | 3 | NZ_ACXX02000001.1 |
| 617 | 14 | 4 | 15 | NZ_JAGE01000001.1 | 19 | 5 | 19 | NZ_ACXX02000002.1 |
| 622 | 4 | 1 | 5 | NZ_JAGE01000001.1 | 5 | 1 | 5 | NZ_ACXX02000002.1 |
| 693 | 6 | 3 | 4 | NZ_JAGE01000001.1 | 5 | 2 | 4 | NZ_ACXX02000002.1 |
| 716 | 3 | 1 | 4 | NZ_JAGE01000001.1 | 5 | 1 | 5 | NZ_ACXX02000002.1 |
| 746 | 6 | 3 | - | - | 3 | 2 | - | - |
| 813 | 7 | 2 | - | - | 8 | 3 | 6 | NZ_ACXX02000007.1 |
| 863 | 5 | 2 | 5 | NZ_JAGE01000002.1 | 5 | 1 | 5 | NZ_ACXX02000003.1 |
| 898 | 7 | 1 | 5 | NZ_JAGE01000002.1 | 7 | 1 | 6 | NZ_ACXX02000003.1 |

| | | | | | | | | |
|---|---|---|---|---|---|---|---|---|
| 915 | 3 | 2 | - | - | - | - | - | - |
| 1000 | 6 | 3 | 6 | NZ_JAGE01000001.1 | 6 | 3 | 6 | NZ_ACXX02000003.1 |
| 1018 | 13 | 1 | 12 | NZ_JAGE01000001.1 | 12 | 0 | 12 | NZ_ACXX02000017.1 |
| 1052 | 4 | 3 | 4 | NZ_JAGE01000001.1 | 3 | 1 | 3 | NZ_ACXX02000017.1 |
| 1073 | 3 | 2 | 3 | NZ_JAGE01000001.1 | 2 | 1 | 2 | NZ_ACXX02000022.1 |
| 1247 | 3 | 1 | 1 | NZ_JAGE01000001.1 | 3 | 0 | 1 | NZ_ACXX02000013.1 |
| 1354 | 3 | 2 | 1 | NZ_JAGE01000001.1 | 1 | 1 | 1 | NZ_ACXX02000010.1 |
| 1382 | 7 | 1 | - | - | - | - | - | - |
| 1435 | 16 | 1 | - | - | - | - | - | - |
| 1745 | 4 | 2 | 6 | NZ_JAGE01000001.1 | 6 | 3 | 6 | NZ_ACXX02000011.1 |
